# Hard-to-beat animacy perception: EEG evidence of accurate animate-inanimate distinction for ambiguous objects

**DOI:** 10.64898/2026.06.15.732334

**Authors:** Farzad Rostami, Céline Spriet, Hans Op de Beeck, Jean-Rémy Hochmann, Liuba Papeo

## Abstract

Human recognition of objects as animate or inanimate is fast and accurate. However, this task may be challenging for objects with animal-like properties (e.g., presence of eyes/face), albeit being inanimate (e.g., a cow-mug). *Lookalike objects* provide an opportunity to examine how visual perception resolves categorical ambiguities and whether it exhibits an intrinsic bias to *see* animacy. During electroencephalography (EEG), we presented healthy humans with images of objects at a regular, rapid frequency (6 Hz), where every five exemplars of a ‘standard’ category (animate or inanimate), an exemplar of the other ‘oddball’ category was shown (1.2 Hz). In some conditions, oddball-stimuli were replaced by lookalikes. Periodic visual stimulation should give rise to a distinctive EEG response at 6 Hz. Moreover, if the oddball category elicits a different neural response than the standard category, a distinct response should be observed at 1.2 Hz. This ‘categorization’ response was found for animate-oddballs among inanimate objects, and *vice versa*. It was also found for lookalike-oddballs presented among animate or inanimate objects, but it was significantly higher in the first case, implying that lookalike objects were perceived as more similar to inanimate than animate objects. The degree of animal resemblance modulated the amplitude of neural response to lookalikes, without however changing the category boundary. These results—also replicated in an artificial model of human vision—demonstrate that animal resemblance of objects is registered in visual perception, but it does not alter the critical ability to distinguish between what is truly alive and what is not.

## Introduction

The property of being—*vs.* not being—alive and animate is a major organizing principle for object representation in visual occipitotemporal cortex: this brain region hosts representational clusters tuned to faces, bodies, body parts and biological motion (Ritchie, et al., 2021; Kanwisher & Yovel, 2006; Downing et al., 2001; Papeo et al., 2017; Pitcher & Ungerleider, 2021); the broad topography of object-related information segregates animate (in ventral aspects) from inanimate (in lateral aspects) objects (Yargholi & Op de Beeck, 2022; Proklova et al., 2019; Konkle & Caramazza, 2013); the distributed patterns of neural activity are more similar between two animates, or between two inanimate objects (Kriegeskorte et al., 2008); and representation of animal species gives rise to a spatial continuum from high-animacy (primates) to low-animacy (bugs) (Connolly et al., 2012; Jozwik et al., 2022; Sha et al., 2015).

Overall, a rich body of work suggests that animate entities may be visually defined by features such as shape, texture, color and spatial frequency spectrum (Jozwik et al., 2016; Long et al., 2017; Nasr et al., 2014; Proklova et al., 2019; Rice et al., 2014; Schmidt et al., 2017; Spriet et al., 2025, Peykarjou et al., 2024), higher-level properties such as, for instance, agency and predictability (Gobbini et al., 2011; Jozwik et al., 2022), and the presence of diagnostic features such as eyes, face and/or body-parts (Bracci et al., 2019; Leys et al., 2025; Proklova & Goodale, 2022; Ritchie et al., 2021).

Under certain spatiotemporal contingencies, individuals can spontaneously attribute animacy to abstract shapes (e.g., triangles and squares) and represent their motion trajectories as parts of meaningful interactions (e.g., triangles chasing a square) (Heider & Simmel, 1944; Wheatley et al., 2007). This suggests that animal appearance is not necessary to represent animacy. However, given certain spatial relations between abstract shapes (e.g., three static dots forming an inverted triangle) or in noisy visual arrays, individuals can *see* faces or bodies, even when there are none (Wardle et al., 2022; Palmer & Clifford, 2020; Rekow et al., 2022). This suggests a particular visual sensitivity to animal appearance. Is animal appearance, that is, the property of having eyes, face and/or body parts, in the absence of other cues of animacy, such as a certain texture, or color or spatial frequency, sufficient to *see animacy*?

Relevant for this fundamental question is the work on the processing of ‘lookalike’ (‘zoomorphic’ is sometimes used synonymously) objects (Bracci et al., 2019; Chen et al., 2023, Duyck et al., 2024). Lookalikes, such as for example a piggybank (i.e., a ceramic coin container in the shape of a pig), are inanimate objects in which inanimate features and animal appearance coexist. That is, they are characterized by presence of eyes, face and/or body-parts, but are made of materials––and therefore texture, colors, glossiness and other features––that are not common in animals, and display features that cue a function or use (in a piggybank, a slot at the top, a rubber plug on the underside). Bracci et al. (2019) showed that human subjects judged lookalikes as more similar to inanimate, than to animate objects, demonstrating to prioritize the (lack of) animacy over animal appearance in explicit categorization. Like humans, artificial deep neural networks (DNNs) modeling human visual processing, classified lookalikes as closer to inanimate than animate objects. However, in specific aspects of the visual cortex, notably in anterior ventral occipitotemporal cortex (VOTC), activity patterns showed the opposite effect: similarity was greater between lookalikes and animates than between lookalikes and inanimate objects. Moreover, activity patterns for lookalikes contained information about their corresponding animal identity but not their corresponding object identity: for example, the pattern for a cow-mug correlated with the pattern for cow but not with the pattern for mug. These findings suggested that animal appearance, conveyed by eyes, faces, and/or body parts, biases object representations toward “animate” in VOTC.

Here, we probed this animal bias in visual object perception, using frequency-tagging electroencephalography (ftEEG), a methodology to measure the immediate, automatic neural response, locked to stimulus appearance, which captures two key aspects of object categorization: discrimination, i.e., the distinct neural responses to stimuli perceived as belonging to different categories, and clustering, i.e., the common neural response to stimuli perceived as belonging to the same category. In ftEEG, a stream of standard stimuli presented at rapid regular stimulation-frequency *Fs* (here, 6 Hz) elicits steady-state visual evoked potentials (SSVP) at the same frequency (Norcia et al., 2015; Retter et al., 2021). If standard stimuli are interleaved by stimuli of a different (oddball) category presented at another regular ‘oddball’ frequency *Fo* (here, 1.2 Hz: 1 oddball every 5 standard stimuli), and if the categorical shift is salient to visual perception, a distinctive response is registered at *Fo*. With this approach, we measured the categorization response to animate objects presented as oddball stimuli in streams of inanimate stimuli (standard stimuli), and *vice versa*. Moreover, we tested the same response to lookalike objects presented as oddballs in streams of either animate or inanimate stimuli.

Building on prior work (Spriet et al., 2025), we expected an oddball response to animates presented among inanimates and *vice versa*. Moreover, we should find an oddball (‘categorization’) response for lookalike-oddballs, if they are perceived as different from animates or inanimates. In particular, if lookalikes are perceived as more similar to animate objects by virtue of their animal-like appearance, the oddball response to lookalikes among inanimate should be comparable to the oddball response to animates among inanimates. Instead, if lookalikes are perceived as more similar to inanimate objects by virtue of the common lack of animacy, the oddball response to lookalikes among animates should be comparable to the oddball response to inanimates among animates.

EEG data analyses were carried out in the frequency and temporal domain. In addition, we tested the representational similarity between the three object categories in the VGG-19 architecture (Simonyan & Zisserman, 2015). Although this convolutional DNN generally reaches human-level performance in object recognition and is used as a model of the primate visual system, animacy represents a case in which object representations in the DNN might not align to those in biological vision (Bracci et al., 2019; Duyck et al., 2024). Thus, the present investigation may also shed light on the conditions (e.g., types of stimuli), under which object representation in an artificial DNN does or does not match representation in biological vision.

To preview, both EEG and DNN results showed that lookalikes were represented as more similar to inanimate than to animate objects. These findings demonstrate that animacy perception is not easily deceived: despite their animal-like appearance, inanimate “lookalike” objects are readily and accurately recognized as such in human vision.

## Materials and Methods

### EEG study

#### Participants

Participants were 21 healthy adults (11 female, mean age = 27.05 years, *SD* = 7.53, range = 18–44 years) with normal or corrected-to-normal vision and no report of psychiatric or neurological conditions. Data from one participant were excluded from analysis as they fell asleep during the experiment, yielding a final sample of 20. An *a priori* power analysis (G*Power 3.1; Faul et al., 2007) was conducted to determine the sample size required to detect an interaction effect comparable to that reported by Rekow et al. (2022), who tested categorical perception of face-like objects with ftEEG and found a significant Category × Duration interaction (η_*p*_^2^ = 0.24). This effect size was converted to Cohen’s 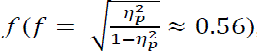, yielding a sample size of 12 to 15 across a plausible range of within-subject correlations (r = 0.3–0.5; α = 0.05; power = 0.95). This experiment was approved by the local ethics committee (CPP Ile de France VIII). All participants provided written informed consent before the experiment and were compensated with money for their time.

#### Stimuli

Stimuli included three sets: animate, inanimate and lookalike objects. The animate and inanimate sets were taken from Spriet et al. (2025). The animate set (n=288) included 154 nonhuman mammals, 68 birds, 46 fish, 14 amphibians, and 6 turtles. Images of humans were excluded to prevent biases elicited by viewing conspecifics (New, Cosmides, Tooby, 2007). Images of insects, arachnids, and reptiles were excluded to prevent reactions such as disgust or fear. The inanimate set (n=288) included 214 artificial objects (17 exemplars of buildings and constructions, 118 exemplars of clothes, pieces of jewelry, buttons, coins, and tools, 39 pieces of furniture and 40 vehicles) and 74 natural objects (49 different fruits and vegetables, and 25 different flowers, bushes and trees). In addition, we created a new set of images featuring 72 lookalike objects, sourced from the Internet. Following Bracci et al. (2019), we considered as “lookalikes”, inanimate objects with both typical animal parts (eyes, faces, bodies and/or body parts) and parts that cue a function or a specific use (e.g., a handle) (Figure 1A).

**Figure 1.**
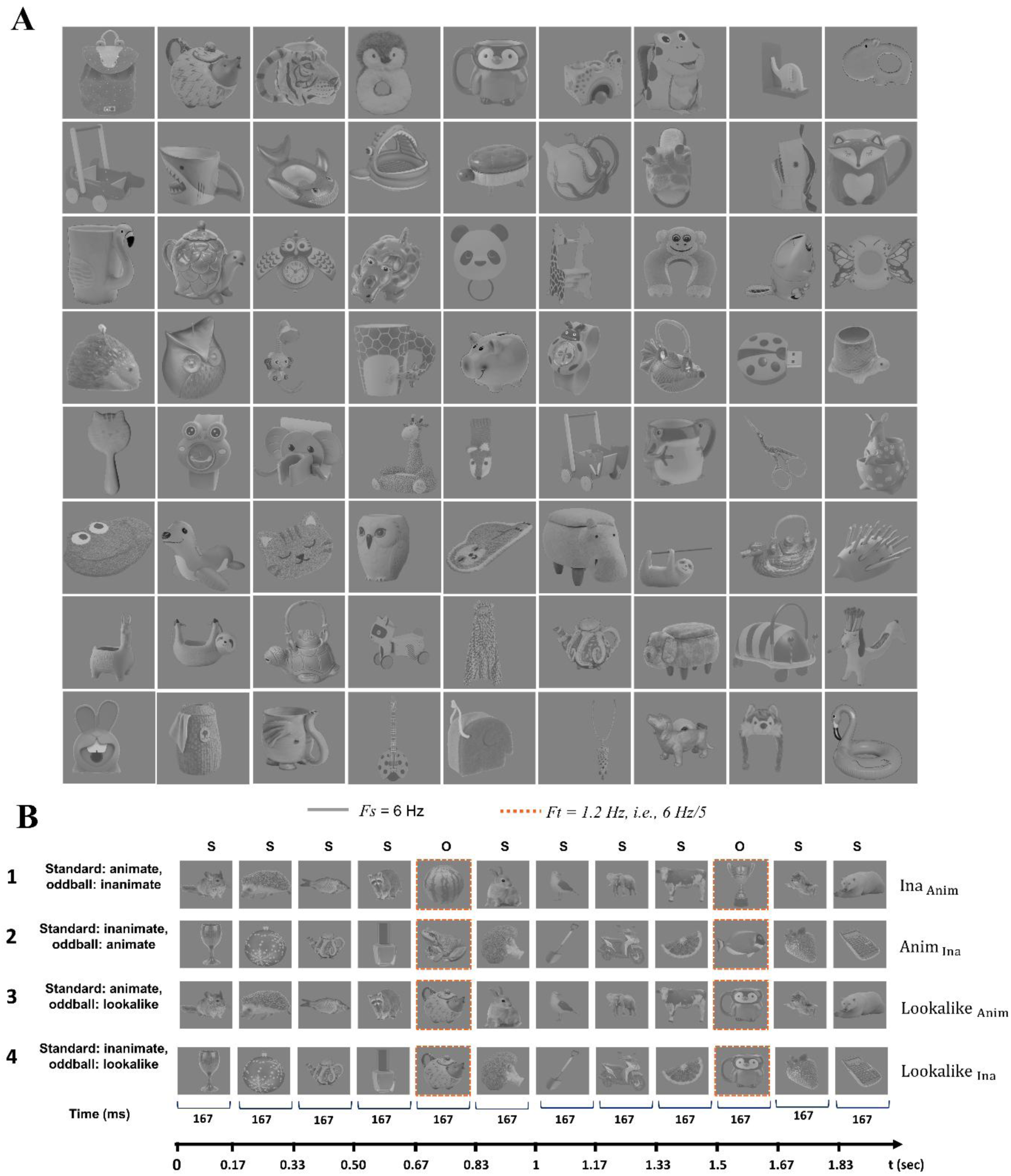
(A) The complete set of 72 lookalike stimuli used in the study. These objects were operationally defined as inanimate items possessing both functional parts (e.g., a handle on a mug) and features characteristic of animals (e.g., eyes, face). (B) Illustration of the experimental stimulation sequences. In each trial, stimuli were presented at a stimulation frequency *Fs* of 6 Hz (6 images per second), with each image displayed for ∼167 ms. An oddball stimulus (squared in orange dotted line) was presented every five standard stimuli (target frequency, *Fo* = 1.2 Hz, i.e., 6 Hz/5). S denotes standard stimuli; O denotes oddball stimuli. The four conditions included in the experiment were: 1) Oddball: inanimate, Standard: animate (*Ina* _*Anim*_); 2) Oddball: animate, Standard: inanimate (*Anim* _*Ina*_); 3) Oddball: lookalike, Standard: animate (*Lookalike* _*Anim*_); and 4) Oddball: lookalike, Standard: inanimate (*Lookalike* _*Ina*_).

We quantified the resemblance of lookalike objects to animals with a rating study involving 20 participants (14 males, 6 females; mean age = 37.35 years, SD = 7.76, range = 23–49 years) external to the main EEG experiment. For data collection, we used the web-based platform Testable (Rezlescu et al., 2020). Participants were presented with 72 grayscale lookalike stimuli, the same used in the EEG experiment. For each stimulus, they rated the degree to which the object resembled an animal on a scale from 0 (no resemblance at all) to 100 (highest resemblance), with responses in steps of 10. Stimuli were presented in random order, and participants completed the ratings without time constraints. Across all 72 lookalike stimuli, the mean animal resemblance rating was 53.66 (SD = 12.75, range = 23.50–83.50), indicating that these objects possessed animal-like features to varying degrees.

Images depicted objects in various size, viewpoint, and lighting. For the EEG study, stimuli were converted to grayscale images, resized to 629 × 629 pixels, and equalized for pixel luminance and root-mean-square contrast using MATLAB (The MathWorks, Natick, MA). With these stimuli, four types of stimulation sequences were created (Figure 1B): 1) Oddball: inanimate, Standard: animate (Ina _Anim_); 2) Oddball: animate, Standard: inanimate (Anim _Ina_); 3) Oddball: lookalike, Standard: animate (Lookalike _Anim_); 4) Oddball: lookalike, Standard: inanimate (Lookalike _Ina_).

#### Procedures

Participants were seated comfortably on a chair approximately 60 cm away from a computer screen (60 Hz refresh rate, resolution 1920 × 1200 pixels, size 51.5 × 32.2 cm), with stimuli presented centrally on the screen, and subtending 16° of visual angle. Before the experiment, participants were instructed to look at the images on the screen without further detail. Stimulus presentation was controlled using MATLAB2015 and PsychToolbox (Brainard, 1997).

In each trial, participants saw one of the four types of sequences with stimuli presented at *Fs* 6 Hz (six images per second, i.e., 166.67 ms per image), a frequency that elicits reliable SSVP in occipitotemporal regions (Rossion et al., 2020). Every five *standard* stimuli (all animates or all inanimate objects), an image of a different category (oddball) was shown, yielding a categorical shift at *Fo* 1.2 Hz. Each trial lasted 60 sec and consisted of a 2-sec fade-in phase, 56 sec of stimulation (288 standards and 72 oddballs), and a 2-sec fade-out phase. In each stimulation sequence, the full sets of standard and oddball stimuli were presented in a randomized order, ensuring that every unique exemplar was shown once per trial. The fade-in and fade-out phases at the beginning and the end of each trial involved gradual changes in image contrast from 0% to 100% (fade-in) and back to 0% (fade-out), to avoid abrupt transitions. Throughout the experiment, each participant saw eight trials of each of four conditions, corresponding to the four types of sequences (Figure 1 B). The 32 trials were grouped in four blocks of eight trials each, two for each condition, with breaks in between. Within each block, the order of conditions was random but two trials of the same condition were always presented consecutively. An orthogonal task was introduced to maintain attention: at the end of each trial, an image was displayed, and participants were asked to indicate whether the image was part of the previous sequence, by pressing a key on a computer keyboard.

#### EEG Recording

EEG data were recorded using a 128-channel Geodesic Sensor Net (GSN; NetStation EGI V2.0) with vertex reference (Cz), continuously digitized at a sampling rate of 1000 Hz using the NetStation acquisition software (EGI V2.0, Net Amp 400 system). Recording took place in a soundproof, dimly lit room to reduce external noise and enhance data quality. Impedance levels were kept as low as possible for each participant and did not exceed 30 kΩ, for optimal signal acquisition. To achieve precise temporal alignment between stimulus presentation and EEG data acquisition, triggers were sent from an experimental computer to the acquisition system via a photosensor positioned at the bottom right of the screen. A white square appeared at this location at the beginning and end of each trial, to precisely mark the timing of each trial and time-lock EEG data to stimulus presentation.

#### EEG Preprocessing

Standard preprocessing was conducted using EEGLAB (Delorme & Makeig, 2004) in MATLAB 2015b. Signals were first filtered using a fourth-order Butterworth bandpass filter with cutoff frequencies of 0.1 Hz and 100 Hz to retain relevant neural oscillations while removing slow drifts and high-frequency noise. For each participant, ‘bad’ channels were identified through visual inspection and interpolated using spherical spline interpolation to maintain the spatial integrity of the electrode montage. Data were then re-referenced to the common average of all 128 electrodes (including interpolated channels), establishing a uniform reference baseline that minimizes bias from any single electrode and enables accurate signal comparison across channels. Independent Component Analysis (ICA) was applied to identify and remove artifacts due to eye movements, blinks, and muscle activity, thereby isolating neural components of interest while preserving the integrity of brain-related signals. After artifact removal, data were segmented into 55-second epochs, each beginning 2.5 seconds after trial onset to exclude the fade-in period and accommodate 66 complete cycles at *Fs* (6 Hz). Finally, trials were averaged separately for each participant for each condition, improving the signal-to-noise ratio by reducing random noise variability and enhancing detection of frequency-tagged responses.

#### EEG Data Analysis

To examine responses in the frequency domain, a Fast Fourier Transform (FFT) was applied to data in each condition after averaging across trials. This yielded a high frequency resolution of 0.018 Hz (1/55 s), enabling precise spectral analysis. The analysis focused on the frequencies of interest, *Fs* (6 Hz) and *Fo* (1.2 Hz), and their harmonics. Baseline-corrected amplitudes at the frequencies of interest were calculated by subtracting the mean amplitude of the 24 surrounding frequency bins (12 on each side, excluding adjacent bins for a range of ±0.48 Hz) from the amplitude of interest, reducing general noise while isolating stimulus-specific neural activity (Retter & Rossion, 2016; Quek & Rossion, 2017; Retter et al., 2021). To define the peaks at the frequencies of interest, the EEG signal of all participants was averaged in the time domain to obtain grand-averaged spectra at each electrode, for each condition. First, the harmonics showing a response were identified following standard procedures (de Heering & Rossion, 2015; Rekow et al., 2022). Second, Z-scores were computed for each harmonic by subtracting the mean amplitude of the local baseline from the amplitude of interest and dividing by the standard deviation of the baseline. For *Fo*, we computed Z-scores for all harmonics below 12 Hz (excluding *Fs* at 6 and 12 Hz). For *Fs*, we computed Z-scores for all harmonics below 50 Hz and selected harmonics with a Z-score higher than 1.64 (p < 0.05, one-tailed). Third, we identified the electrodes showing a significant response at the selected harmonics. This was done by computing a Z-score on the grand-averaged spectrum for each electrode independently. Electrodes were selected if the corresponding Z-score exceeded 3.33 (corresponding to *p* = 0.0004, one-tailed, Bonferroni-corrected for 128 electrodes).

Statistical analyses were conducted using parametric tests on baseline-corrected amplitudes. In particular, we assessed whether the amplitude of the response at *Fs* and *Fo* was significantly higher than the noise level. To this end, each participant’s baseline-corrected amplitude, summed over the selected harmonics and averaged over the selected electrodes, was tested against 0 (*t* test, one-tail). Differences between conditions and their spatial distribution were assessed with a cluster-based permutation analysis correcting for multiple comparisons. In this analysis, we used a 2 × 2 repeated-measures ANOVA with within-subject factors Standard (Animate vs. Inanimate) and Oddball (Lookalike vs. Other –where ‘other’ could be either the animate or the inanimate objects used as oddballs). The analysis was performed across all 128 electrodes using baseline-corrected amplitudes summed across *Fo* and its first 8 harmonics (1.2 Hz to 9.6 Hz). Crucially, the same set of harmonics was used for all conditions to ensure unbiased comparison. Significant clusters included a minimum of 2 neighboring electrodes identified with 5,000 permutations and a cluster-forming threshold of α = 0.05. For each permutation, condition labels were randomly shuffled, and cluster-level statistics were computed. The significance probability of the observed clusters was determined by comparing the cluster-mass statistics to the null distribution generated from permuted data. This approach controls the family-wise error rate while maintaining sensitivity to spatially contiguous effects (Maris & Oostenveld, 2007).

To test the interaction within each cluster, pairwise *t* tests between experimental conditions (Ina _Anim_, Anim _Ina_, Lookalike _Anim_, and Lookalike _Ina_) were performed on the baseline-corrected amplitudes extracted from the cluster (α = 0.0125, Bonferroni-corrected for the number of comparisons).

#### ERP Preprocessing

To investigate the latencies of possible categorization effects, we conducted ERP analysis on preprocessed EEG data. EEG data were segmented into short epochs for each of experimental condition, separately. Each epoch was time-locked to the onset of an oddball stimulus within an “SOSSS” sequence, where “S” stands for standard stimulus “O” stands for oddball stimulus. Each epoch spanned 833 ms, corresponding to five stimulus presentations at *Fs* of 6 Hz (5 × 166.67 ms). This approach allowed us to examine the temporal dynamics characterizing to the categorical shift at each oddball stimulus. Baseline correction was applied using the pre-stimulus interval (−167 to 0 ms), corresponding to the duration of the standard stimulus immediately preceding the oddball. Epochs with excessive noise or artifacts, primarily from eye blinks and muscle movements, were excluded using both automated algorithms and visual inspection to maintain data quality. Next, a low-pass filter with a 30 Hz cutoff was applied to each epoch, reducing high-frequency noise and preserving relevant ERP components. A multi-notch filter was applied to remove the *Fs* (6 Hz) and harmonics, thus isolating neural responses associated with the categorical shift at *Fo* (1.2 Hz).

#### ERP Data Analysis

ERP data were averaged across the electrodes forming the clusters that showed an interaction effect in the frequency-domain analysis (see EEG Data Analysis). For each cluster, a cluster-based permutation analysis was used to test the Standard (Animate vs. Inanimate) × Oddball (Lookalike vs. Other) interaction across the epoch to identify temporal windows where the interaction was significant. Neighboring time points with a *t* value exceeding a predefined threshold (corresponding to p < .05) were grouped into clusters. Statistical significance was assessed with cluster-mass permutation based on *t* tests with 5,000 iterations, in which condition labels were randomly shuffled, and cluster-level statistics were recomputed for each permutation (Maris & Oostenveld, 2007). The significance of each observed cluster was determined by the proportion of permutations yielding a summed *t* value higher than the observed cluster statistic. Paired-samples *t* tests were then conducted on mean amplitudes extracted from the interaction-defined temporal window(s), to characterize the source of the interaction. This two-step procedure was applied in both the central and the posterior cluster.

#### Artificial Deep Neural Network

We measured the representational similarity between animate, inanimate, and lookalike objects using the VGG-19 architecture (Simonyan & Zisserman, 2015). VGG-19 is a convolutional deep neural network (DNN) used as a model of the primate visual system, particularly the ventral occipitotemporal cortex (VOTC), which achieves human-level object recognition performance through a series of processing stages (layers) that, in a feedforward manner, nonlinearly transform an input image volume into a one-dimensional vector containing the class scores. We used the pretrained version (MatConvNet; Vedaldi & Lenc, 2015), trained on ∼1.2 million images from the ImageNet database (ILSVRC2012) encompassing 1,000 object categories including animals (40%) and inanimate objects (60%). Before analysis, standardized preprocessing steps included mean subtraction of the training images and resizing of all stimuli to 224 × 224 pixels.

Using the Deep Learning Toolbox in MATLAB (MathWorks, 2024), for each of our stimuli, we extracted feature representations from the final fully connected layer (fc8) of VGG-19, the best candidate for VOTC representations (Khaligh-Razavi & Kriegeskorte, 2014; Cichy et al., 2016). To quantify the representational dissimilarity between stimuli, we computed a representational dissimilarity matrix (RDM; Kriegeskorte et al., 2008), in which each cell represented the dissimilarity between each pair of stimuli as 1 – *r*, where *r* is the Pearson correlation coefficient between the feature vectors of the two stimuli in the fc8 layer. This correlation-based distance metric provided a measure of representational divergence, with higher values indicating greater dissimilarity between stimuli. To determine whether lookalike objects were represented more similarly to animate or inanimate objects in VGG-19, we implemented the following steps: For each of the 72 lookalikes, we computed the correlation with each of the 288 animate stimuli and averaged the 288 correlation coefficients to obtain the mean similarity to animates. We repeated this procedure with the 288 inanimate stimuli to obtain the mean similarity to inanimates for each lookalike. This yielded two paired distributions of 72 values. We then used a paired-sample *t*-test to compare these two distributions, testing whether lookalike stimuli were, on average, represented more similarly to the animate or inanimate category.

## Results

### Frequency-tagging EEG

We tested whether the rapid presentation of a sequence of images elicited an EEG response at the frequency *Fo* of the oddball category, signaling automatic detection of the categorical shift. We expected to observe this response for oddball-animates presented among standard-inanimate, and *vice versa*. In addition, we measured and compared the effect for lookalikes when presented as oddball stimuli among animate or inanimate objects. The categorical discrimination denoted by the amplitude of the oddball response will indicate whether lookalikes are represented as more similar to animates or to inanimate objects. Lower discrimination (i.e., a smaller oddball response) for lookalikes presented among animates would suggest that lookalikes are visually categorized mainly based on their animal-like appearance (which they shared with animates). Lower discrimination (i.e., a smaller oddball response) for lookalikes presented among inanimates would suggest that lookalikes are visually categorized mainly based on their *true* inanimacy (which they shared with inanimates).

First, a response at *Fs* (6 Hz) and harmonics indicated reliable synchronization of neural activity with visual stimulation. All harmonics below 50 Hz showed a significant response (*z* > 1.64) widely distributed over the scalp, in all four experimental conditions (Ina _Anim_: *M*_*Amiplitude*_±*SD* = 0.378±0.120; 95% *CI* = 0.322 – 0.434; t(19) = 14.061; *p* < 0.0001; *d* = 3.144; Anim _Ina_: *M*_*Amiplitude*_±*SD* = 0.393±0.116; 95% *CI* = 0.339 – 0.447; t(19) = 15.201; p < 0.0001; d = 3.399; Lookalike _Anim_: *M*_*Amiplitude*_±*SD* = 0.384±0.124; 95% *CI* = 0.326 – 0.442; *t*(19) = 13.863; *p* < 0.0001; *d* = 3.100; Lookalike _Ina_: *M*_*Amiplitude*_±*SD* = 0.359±0.124; 95% *CI* = 0.301 – 0.417; *t*(19) = 12.995; *p* < 0.0001; *d* = 2.906).

Next, we analyzed the responses at *Fo* (1.2 Hz) and harmonics, as an index of visual categorization (Figure 2A). A significant (above noise-level) response, peaking over posterior electrodes, was found in all four conditions (Ina _Anim_: first six harmonics of *Fo*; *M*_*Amplitude*_±*SD* = 0.354±0.159; 95% *CI* = 0.280 – 0.428; *t*(19) = 9.982; *p* < 0.0001; *d* = 2.232; Anim _Ina_: first four harmonics; *M*_*Amplitude*_±*SD* = 0.183±0.091; 95% *CI* = 0.140 – 0.225; *t*(19) = 9.029; *p* < 0.0001; *d* = 2.019; Lookalike _Anim_: first four harmonics; *M*_*Amplitude*_±*SD* = 0.127±0.066; 95% *CI* = 0.096 – 0.157; *t*(19) = 8.567; *p* < 0.0001; *d* = 1.916; Lookalike _Ina_: first five harmonics; *M*_*Amplitude*_±*SD* = 0.285±0.147; 95% *CI* = 0.216 – 0.354; *t*(19) = 8.657; *p* < 0.0001; *d* = 1.936). These results showed successful categorization in all four conditions. That is, participants accurately discriminated between animate, inanimate and lookalike objects.

**Figure 2.**
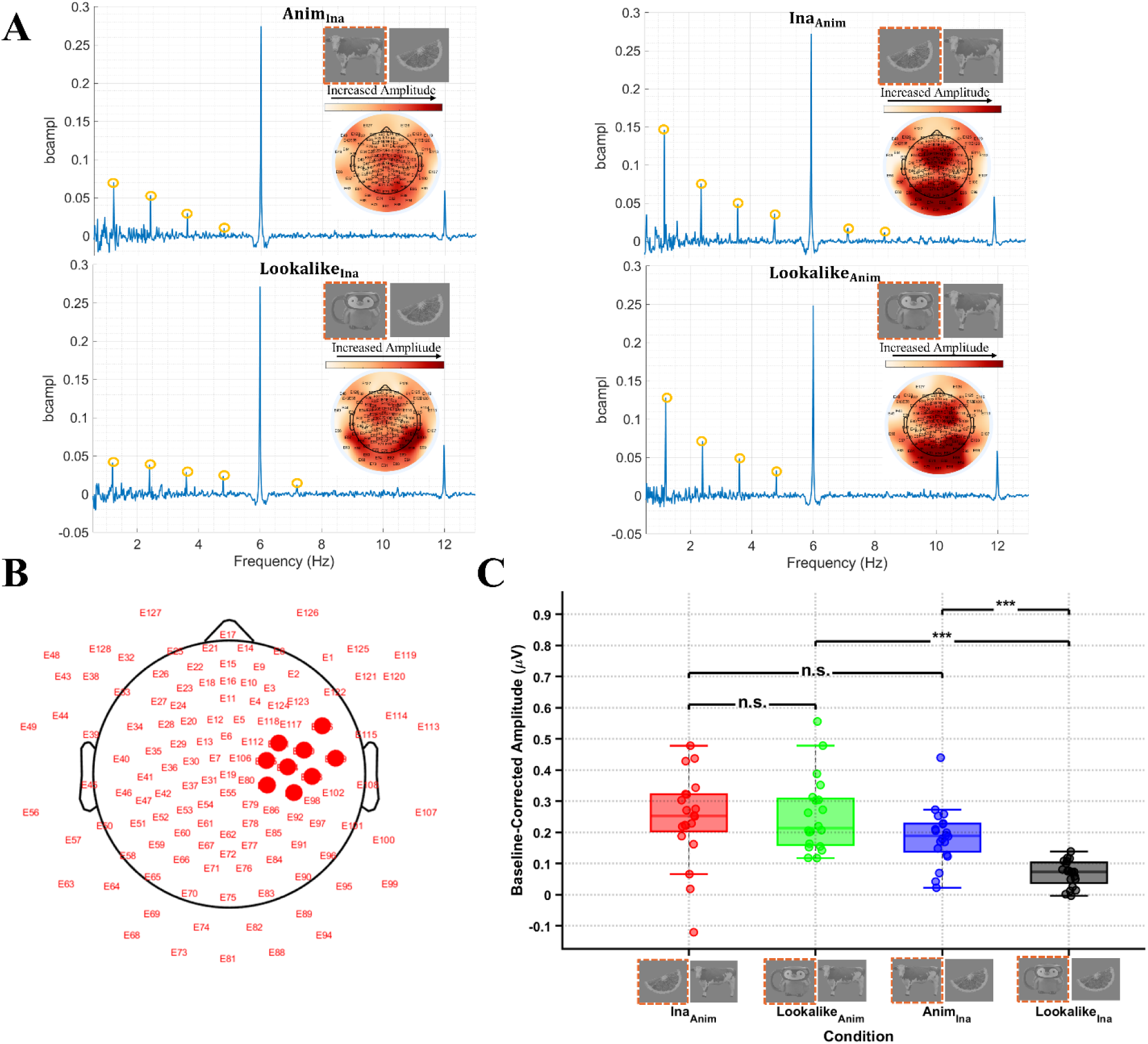
(A) Grand-averaged baseline-corrected amplitude spectra at the oddball frequency *Fo* (1.2 Hz) and its harmonics. Yellow circles highlight harmonics showing significant responses above noise level in each condition. Statistical comparisons across conditions were performed using the same set of harmonics (first 8 harmonics: 1.2–9.6 Hz) for all conditions. Topoplots denote the scalp distribution of the effects. (B) Topographical map of the cluster showing the interaction between Standard and Oddball (electrodes E87, E93, E103, E104, E105, E109, E110, E111). (C) Baseline-corrected amplitudes extracted from the cluster in B for the four experimental conditions. Error bars represent SEM; *** *p* < 0.001; n.s. = not significant.

To test the differences between conditions, a cluster-based permutation analysis tested a 2 Standard (Animate vs. Inanimate) × 2 Oddball (Lookalike vs. Other –where ‘other’ could be either the animate or the inanimate objects used as oddballs) interaction across all 128 electrodes using baseline-corrected amplitudes summed across the first 8 harmonics of *Fo* (1.2 Hz to 9.6 Hz) for all conditions. A main effect of Standard was identified in a widespread cluster of 90 electrodes (cluster-mass stat = 487.32, *p* < 0.001). This effect indicated that the oddball response was overall stronger when standard stimuli were animate (*vs* inanimate) objects. There was also a main effect of Oddball in a cluster of 33 electrodes distributed over occipital to frontal regions (cluster-mass stat = 156.84, *p* = 0.008), showing that the oddball response was overall weaker when oddball were lookalikes (*vs* ‘others’, i.e., either animate or inanimate). Finally, a significant interaction between Standard and Oddball was found in a cluster of 8 electrodes over right central regions (cluster-mass stat = 42.67, p = 0.019) (Figure 2B).

Pairwise *t* tests performed over the baseline-corrected amplitudes extracted from this ‘interaction’ cluster showed that the effect of oddball-lookalikes was stronger when they appeared among animate than among inanimate objects (Lookalike _Anim_ vs. Lookalike _Ina_: *M*_*Difference*_±*SD* = 0.188±0.117; 98.75% *CI* = [0.116, 0.261]; *t*(19) = 7.205, *p* < 0.001, *d* = 1.611). In effect, in the first of the two conditions (oddball-lookalikes among animates), the oddball effect was comparable to the effect observed when inanimate objects appeared as oddball stimuli among animates (Ina _Anim_ vs. Lookalike _Anim_: *M*_*Difference*_±*SD* = –0.011±0.118; 98.75% *CI* = [-0.084, 0.062]; *t*(19) = –0.418, *p* = 1.000, *d* = –0.093). That is, this cluster discriminated between animates and inanimates as well as between animates and lookalikes. In the other condition (oddball-lookalikes among inanimate objects), the oddball effect was smaller than the effect observed when animate objects appeared as oddball stimuli among inanimate objects (Anim _Ina_ vs. Lookalike _Ina_: *M*_*Difference*_±*SD* = 0.119±0.103; 98.75% *CI* = [0.056, 0.182]; *t*(19) = 5.187, *p* < 0.001, *d* = 1.160). In sum, this cluster discriminated between inanimate and animate objects better than between inanimate objects and lookalikes. The oddball effect was comparable between conditions Ina _Anim_ and Anim _Ina_ (*M*_*Difference*_±*SD* = 0.059±0.107; 98.75% *CI* = [-0.007, 0.124]; *t*(19) = 2.454, *p* = 0.096, *d* = 0.549). Note that because the cluster for this analysis was selected based on a previous cluster-mass analysis looking for a significant interaction, *t* tests served to characterize the nature of the effect rather than provide independent hypothesis testing.

### ERPs

An ERP analysis was carried out to assess effects found in the above central cluster of electrodes, in the temporal domain. For each condition, we averaged the ERP data across the 8 central electrodes forming the cluster showing a significant Standard × Oddball interaction in the frequency-domain analysis (Figure 2B), and used cluster-mass permutation analysis to identify the temporal windows in which the interaction was significant. Note that because this cluster was selected based on electrodes showing a significant interaction in the frequency-domain analysis, the following ERP analyses served to characterize the temporal dynamics of the effect rather than provide independent statistical testing. A cluster-based permutation test on the Standard × Oddball interaction across the epoch identified a temporal window between 411 and 677 ms post-stimulus onset (cluster-level *p* < 0.01). Pairwise *t* tests on mean amplitudes extracted from this window revealed that, in the context of inanimate standards, the ERP response to animate oddballs differed significantly from the response to lookalike oddballs, *t(19)* = −3.645, *p* = 0.002, *M_difference_* ±*SD* = −0.255±0.313 µV, *d* = −0.815. In contrast, in the context of animate standards, the ERP response to inanimate oddballs was comparable to the response to lookalike oddballs, *t(19)* = −0.873, *p* = 0.394, *M_difference_* ±*SD* = −0.067±0.341 µV, *d* = −0.195.

These results indicate that, when inanimate objects served as standards, animate oddballs were discriminated more strongly than lookalike oddballs, whereas when animate objects served as standards, inanimate and lookalike oddballs evoked comparable responses. These ERP effects in the cluster of central electrodes are consistent with the frequency-tagging effects found in the same cluster, showing that in streams of animate-standards, lookalikes were discriminated as well as inanimate objects, whereas in streams in inanimate-standards, animates yielded greater oddball responses (i.e., denoting stronger discrimination) than lookalike objects (Figure 2C).

Additional analyses showed that, in this central cluster, although lookalikes were represented as more similar to inanimate than animate objects, the degree of animal-like appearance of lookalikes modulated the oddball-ERP amplitude. In particular, we considered the ERPs extracted from the central interaction-cluster in a time window of 400–600 ms. We found that, for lookalikes presented among inanimate objects, the ERP amplitude of the oddball response was greater the higher the animal-like appearance, as measured with the human ratings (see Stimuli) (*r* = 0.526, *p* = 0.012; Figure 3B, right). Instead, for lookalikes presented among animate objects, the ERP amplitude was lower the higher the animal-like appearance (*r* = –0.124, *p* = 0.042; Figure 3B, left). These correlations suggest that the animal-like appearance modulated neural response to lookalike objects in a way that depended on the context: among inanimate objects, lookalikes *looked odder* (greater oddball ERP amplitude) the more they looked like animals (higher animal-like appearance); among animate objects, lookalikes *looked odder* (greater oddball ERP amplitude) the less they looked like animals (lower animal-like appearance).

**Figure 3.**
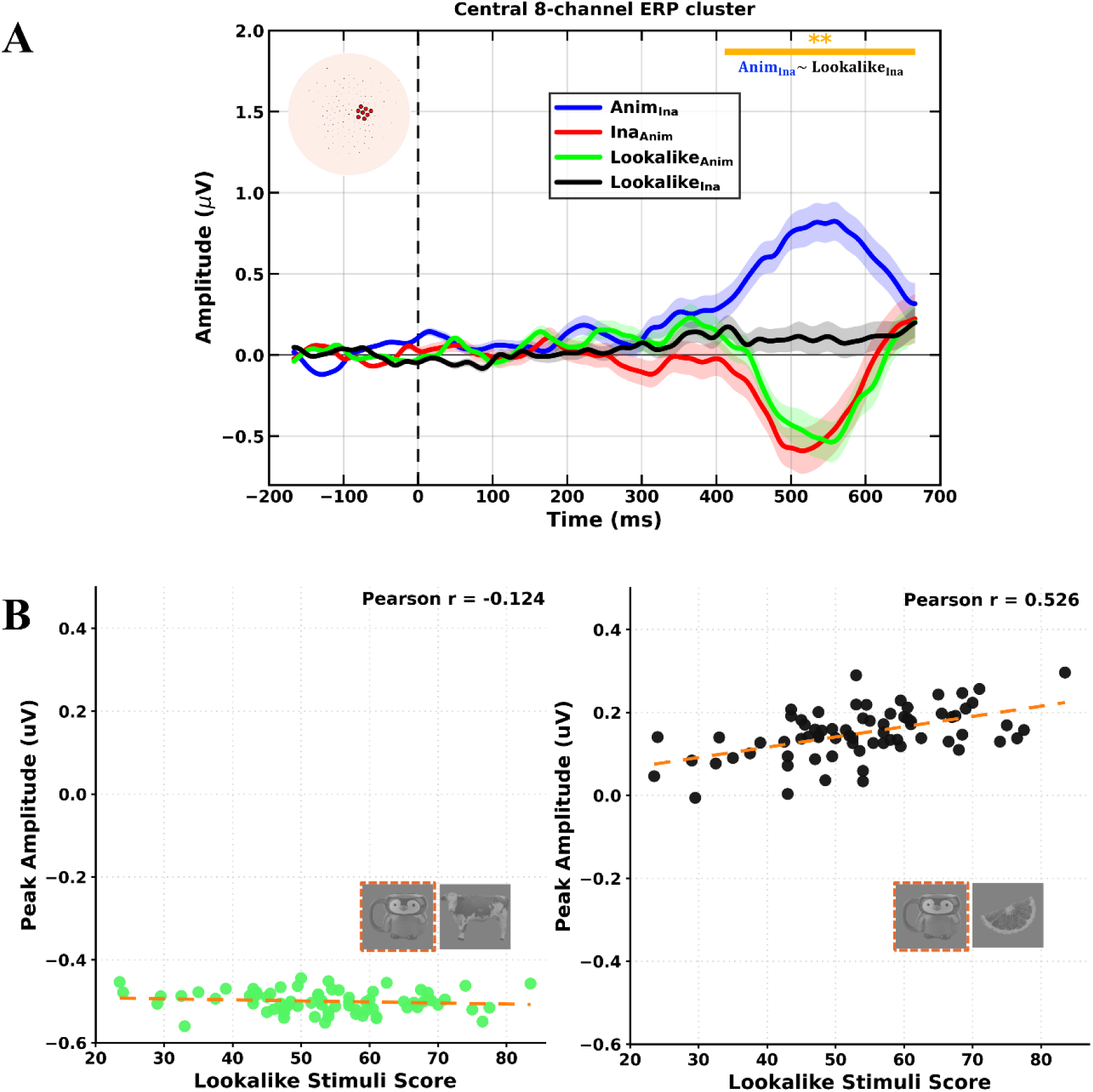
Central-cluster ERP effects and modulation by animal-like appearance. (A) Grand-averaged ERP waveforms (N = 20) over the 8-electrode central cluster (E87, E93, E103, E104, E105, E109, E110, E111) identified by the frequency-domain Standard × Oddball interaction, after removal of Fs and its harmonics. Shaded areas denote ± SEM. Cluster-based permutation testing revealed a significant Standard × Oddball interaction between 411 and 677 ms post-stimulus onset (cluster-level p < 0.01). Within this window, ERPs to animate and lookalike oddballs differed significantly in the context of inanimate standards (Anim _Ina_ vs. Lookalike _Ina_), whereas inanimate and lookalike oddballs did not differ in the context of animate standards (Ina _Anim_ vs. Lookalike _Anim_). (B) Correlation between behavioral ratings of animal-like appearance (0–100 scale) and oddball-ERP amplitude for lookalike stimuli in the central cluster (400–600 ms). Left: Lookalike _Anim_ condition (green; r = −0.124, p = 0.042). Right: Lookalike _Ina_ condition (black; r = 0.526, p = 0.012). Each point represents one of the 72 lookalike stimuli.

Given the correlations between degree of animal-like appearance and oddball-ERP responses, we tested whether the ERP effects illustrated in Figure 3A changed, when only lookalikes with the highest animal-like appearance (the top 25% based on ratings) were considered. To this end, we divided the lookalikes into two sets based on the independent ratings: high animal-like appearance (top 25%, mean rating = 69.81 ± 5.64) and low animal-like appearance (bottom 25%, mean rating = 37.61 ± 7.41). Results showed that high animal-like appearance did not change the category boundaries in the central interaction-cluster: the oddball-ERP amplitude for lookalikes with high animal-like appearance presented among inanimates was still smaller than the same response to animates among inanimates (Anim _Ina_ vs. Lookalike _Ina__H: *M*_*Difference*_±*SD* = 0.19±0.11 µV; 98.75% *CI* = [0.09, 0.29]; temporal cluster: 420-630 ms; cluster-mass permutation *t t*ests: cluster-level *p* < 0.05); and the oddball-ERP amplitude for lookalikes with high animal-like appearance presented among animates was still comparable to the same response measured for inanimates among animates (Ina _Anim_vs. Lookalike _Anim__H: *M*_*Difference*_±*SD* = 0.03±0.09 µV; 98.75% *CI* = [-0.05, 0.11]; cluster-mass permutation t tests: *p* > 0.05). In other words, in the central cluster, animate objects were discriminated from inanimate objects better than lookalike objects, even when lookalikes had high animal resemblance. Instead, inanimate objects were discriminated from animate objects as well as lookalike objects, even when lookalikes had high animal resemblance. The same analyses were repeated considering lookalikes with the lowest animal resemblance, which yielded the same results (Anim _Ina_ vs. Lookalike _Ina__L: *M*_*Difference*_±*SD* = 0.21±0.10 µV; 98.75% *CI* = [0.12, 0.30]; cluster-level *p* < 0.05; Ina _Anim_ vs. Lookalike _Anim__L: *M*_*Difference*_±*SD* = 0.02±0.08 µV; 98.75% *CI* = [-0.05, 0.09]; *p* > 0.05).

### Analyses in posterior (visual) areas

In prior fMRI studies (Bracci et al., 2019), the *animacy effect* of lookalikes—i.e., a greater representational similarity between lookalike and animate (vs. inanimate) objects—was specifically observed in posterior occipital-temporal cortex. The above interaction, showing greater representational similarity between lookalike and inanimate objects, was found in a cluster of electrodes with central distribution. The response in posterior electrodes (visual areas) remains unclear. To address this, we conducted an exploratory ERP analysis over a posterior occipitotemporal electrode set.

We analyzed ERPs over a set of 33 occipitotemporal electrodes (E58, E59, E60, E61, E62, E63, E64, E65, E66, E67, E69, E70, E71, E72, E73, E74, E75, E76, E77, E78, E81, E82, E83, E84, E85, E88, E89, E90, E91, E94, E95, E96, E99), selected to sample posterior visual scalp regions. A cluster-based permutation test of the Standard (Animate, Inanimate) × Oddball (Lookalike, Other) interaction identified two significant temporal windows: an early window from 227 to 349 ms post-stimulus onset (cluster-level *p* < 0.05) and a later window from 461 to 665 ms post-stimulus onset (cluster-level *p* < 0.01) (Figure 4A). Follow-up pairwise comparisons on mean amplitudes extracted from each window were used to characterize this interaction (all *p* values corrected for multiple comparisons).

**Figure 4.**
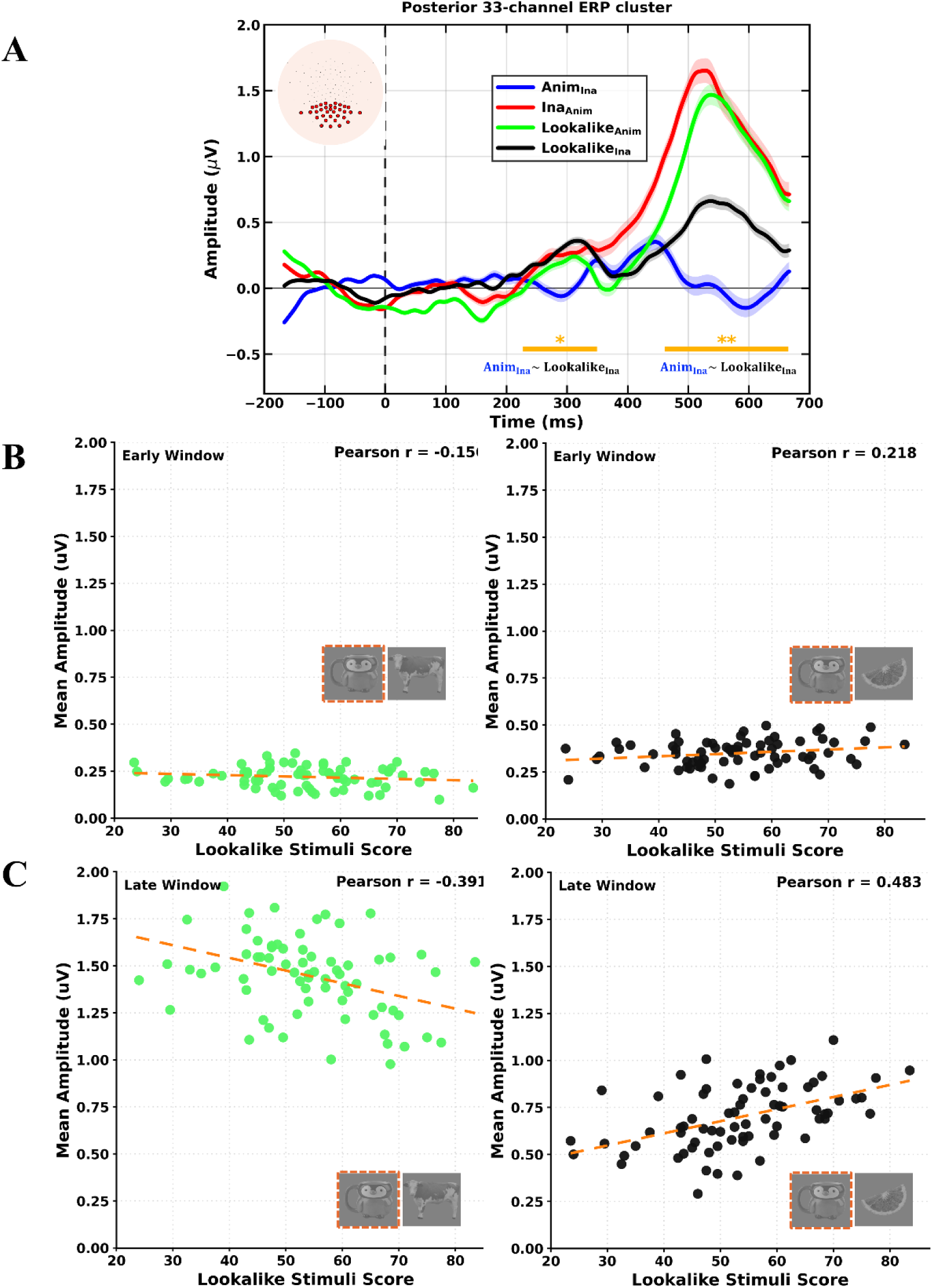
Posterior occipitotemporal ERP effects and modulation by animal-like appearance. (A) Grand-averaged ERP waveforms (N = 20) over the 33-electrode posterior occipitotemporal ROI (E58, E59, E60, E61, E62, E63, E64, E65, E66, E67, E69, E70, E71, E72, E73, E74, E75, E76, E77, E78, E81, E82, E83, E84, E85, E88, E89, E90, E91, E94, E95, E96, E99) used for the hypothesis-driven ERP analysis. Shaded areas denote ± SEM. Cluster-based permutation testing identified two significant temporal windows for the Standard × Oddball interaction: an early window from 227 to 349 ms post-stimulus onset (cluster-level p < 0.05) and a later window from 461 to 665 ms (cluster-level p < 0.01). In both windows, ERPs to animate and lookalike oddballs differed in the context of inanimate standards (Anim _Ina_ vs. Lookalike _Ina_), whereas inanimate and lookalike oddballs did not differ in the context of animate standards (Ina _Anim_ vs. Lookalike _Anim_). (B) Correlation between behavioral ratings of animal-like appearance (0–100 scale) and oddball-ERP amplitude for lookalike stimuli in the early posterior window (227–349 ms). Left: Lookalike _Anim_ condition (green; r = −0.156, p = 0.231). Right: Lookalike _Ina_ condition (black; r = 0.218, p = 0.089). (C) Same as in (B), for the later posterior window (461–665 ms). Left: Lookalike _Anim_ condition (green; r = −0.391, p = 0.048). Right: Lookalike _Ina_ condition (black; r = 0.483, p = 0.021). In both panels (B) and (C), each point represents one of the 72 lookalike stimuli.

In the early window, significantly different ERP response was found for animate oddballs and lookalike oddballs presented in the context of inanimate standards, *t(19)* = 2.152, *p* = 0.045, *M_difference_* ±*SD* = 0.241 ± 0.502 µV, *d* = 0.482. By contrast, no difference was found between inanimate oddballs and lookalike oddballs in the context of animate standards, *t(19)* = −0.588, *p* = 0.563, *M_difference_* ±*SD* = −0.080 ± 0.608 µV, *d* = −0.131. The later window showed the same pattern. Animate and lookalike oddballs again differed significantly in the context of inanimate standards, *t(19)* = 3.208, *p* = 0.0046, *M_difference_* ±*SD* = 0.499 ± 0.696 µV, *d* = 0.717, whereas responses were comparable for inanimate and lookalike oddballs in the context of animate standards, *t(19)* = −1.385, *p* = 0.182, *M_difference_* ±*SD* = −0.155 ± 0.500 µV, *d* = −0.310.

These results show context-dependent differences in posterior electrodes in two temporal windows (Figure 4A). In particular, there was a reliable difference between animate and lookalike oddballs presented in the inanimate-standard context, but no difference between inanimate and lookalike objects presented in the animate-standard context. However, this posterior effect (Figure 4A) differed from the effect observed in the central cluster (Figure 3A) in both latency and polarity. It had an earlier onset (from 227 ms vs. 411 ms post-stimulus onset), and the direction of the pairwise difference was reversed at the scalp level: in the posterior cluster, the response to animate oddballs was smaller than the response to lookalike oddballs in the context of inanimate standards, whereas in the central cluster the response to animate oddballs was greater. Because scalp ERP polarity depends on the location and orientation of the underlying generators, an opposite waveform direction across posterior and central electrodes does not by itself imply an opposite underlying categorical effect. However, given differences in polarity and latency, we would not argue that the two clusters captured the same processing but that they converged in showing quantitatively similar effects of lookalikes and inanimates, and more dissimilar effects of lookalikes and animates.

Finally, we examined whether ERP response to lookalikes in the posterior cluster covaried with the degree of animal-like appearance. Correlation analyses were carried out separately for the two temporal windows (early: 227–349 ms; late: 461–665 ms). For each window, peak ERP amplitudes in response to lookalikes were extracted and correlated with ratings measuring the lookalikes’ animal-like appearance. Results were compatible with the pattern observed in the central cluster and illustrated in Figure 3B. In particular, in the early window, for lookalikes presented among inanimates, the ERP amplitude increased for lookalikes with high resemblance to animals, although this only emerged as a trend, *r* = 0.218, *p* = 0.089 (Figure 4B, right). For lookalikes presented among animate objects, the ERP amplitude decreased for lookalikes with high resemblance to animals, although this correlation was not significant, *r* = −0.156, *p* = 0.231 (Figure 4B, left). Correlations showed the same pattern and were statistically significant, considering the responses in the later window (461–665 ms). For lookalikes presented among inanimate objects, peak ERP amplitudes increased with the increase in animal-like appearance, *r* = 0.483, *p* = 0.021 (Figure 4C, right). For lookalikes presented among animate objects, peak ERP amplitude decreased with the increase in animal-like appearance, *r* = −0.391, *p* = 0.048 (Figure 4C, left).

### Artificial Deep Neural Network

We measured the representational similarity between animate, inanimate, and lookalike objects within the VGG-19 (layer fc8), considering the correlations between lookalikes and animates and lookalikes and inanimates. Results are displayed in Figure 5. Each entry in the RDM represents the dissimilarity between a pair of stimuli, with higher values indicating greater representational divergence. The matrix revealed distinct clustering patterns corresponding to the stimulus categories. The distribution of correlation values between lookalikes and inanimates had a mean of *r* = 0.62 (*SD* = 0.05), indicating relatively high representational similarity between the two categories. The distribution of correlation values between lookalikes and animates had a mean of *r* = 0.28 (*SD* = 0.07), indicating lower similarity. Statistical comparison between the two distributions confirmed that representation of lookalike stimuli in the VGG-19 correlated more–denoting higher similarity–with representation of inanimate objects than animate ones, *t*(71) = 10.35, *p* < 0.001, Cohen’s d = 1.22.

**Figure 5.**
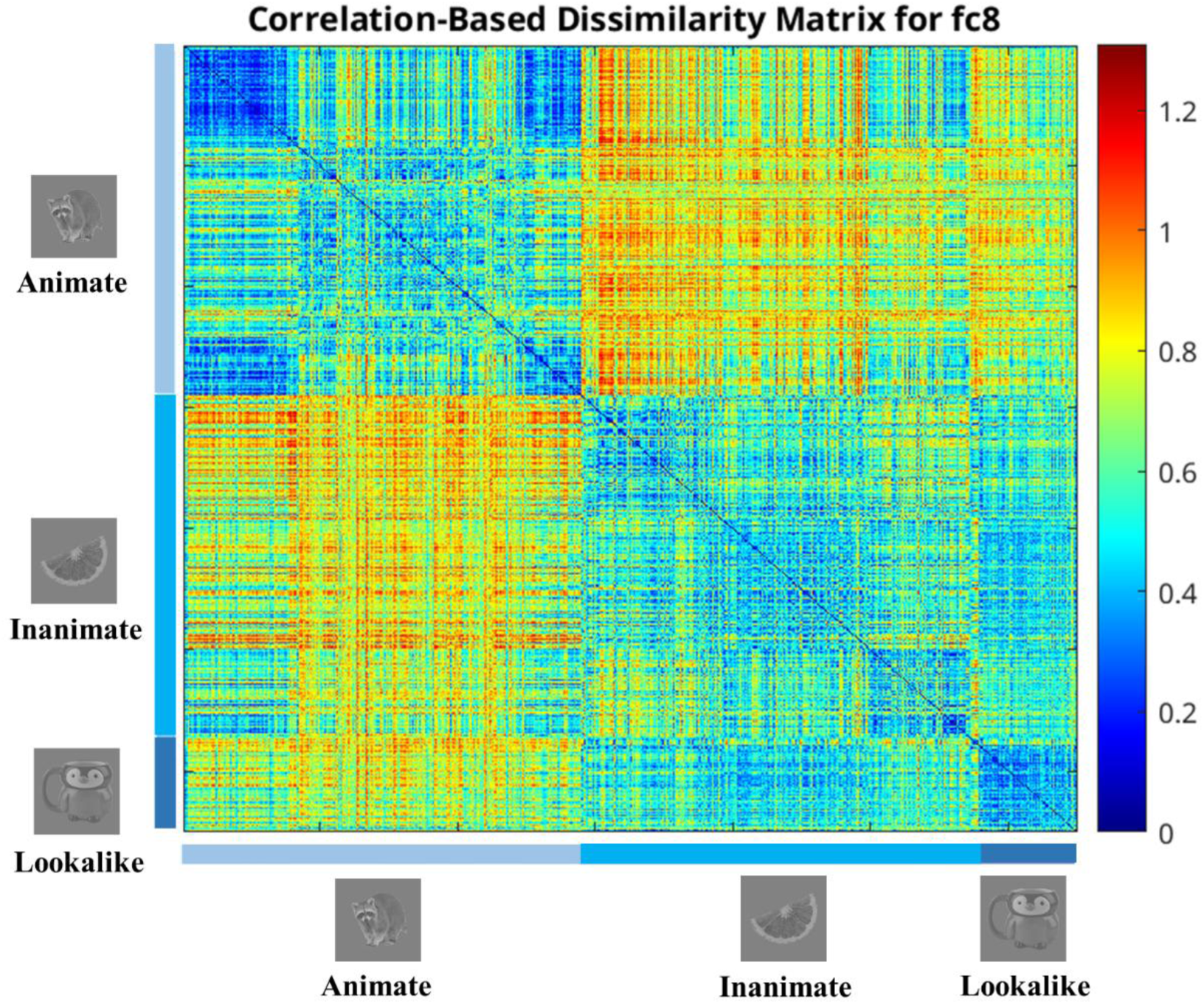
Correlation-Based Dissimilarity Matrix for the fully connected eighth layer (fc8) of the VGG19 network. The matrix illustrates the representational dissimilarity between ‘Animate’, ‘Inanimate’, and ‘Lookalike’ images. The dissimilarity between ‘Lookalike’ and ‘Inanimate’ images is less than that between ‘Lookalike’ and ‘Animate’ images, indicating that lookalike images are represented more similarly to inanimate objects within the network.

These findings indicate that, within the representational space of VGG-19, lookalike objects are represented as more similar to inanimate objects than to animate ones. This pattern aligns with the EEG effects suggesting greater representational distinctiveness for animate objects and greater similarity of the responses to inanimate and lookalike objects. Given that VGG-19 is commonly interpreted as a computational model of ventral visual stream processing, particularly occipitotemporal cortex, the alignment between the DNN’s representational space and our EEG findings underscores the potential of DNNs as models for understanding how the human visual system achieves categorical object recognition.

## Discussion

Object categorization by animacy has been proposed as a major organizing principle of object representation in visual cortex (Grill-Spector and Weiner, 2014), the first to emerge in infancy (Spriet et al., 2022), and one of the most efficient mechanisms of visual perception (Thorpe et al., 1996; VanRullen and Thorpe, 2001). Yet, the features that are necessary and sufficient to *see* animacy remain debated. Here we investigated whether vision discriminates *looking like an animal* from truly *being an animal*––i.e., whether the degree of animal appearance trumps the representation of animacy.

We used EEG measures of neural discrimination between animate and inanimate objects, and studied how these two categories were represented with respect to *lookalike*, visually ambiguous objects. We reasoned that visual object categorization based on animal-like appearance should increase the representational similarity between lookalike and animate objects, yielding comparable oddball response (i.e., our measure of discrimination) to animates among inanimates and lookalikes among inanimates. Instead, visual categorization by true animacy should results in greater representational similarity between lookalike and inanimate objects, yielding comparable oddball response to inanimates among animates and lookalikes among animates.

Results of the EEG responses at the oddball frequency supported the latter prediction: lookalike objects were perceived as more similar to inanimate than animate objects. Complementary ERP analyses showed that a pattern denoting greater representational similarity between lookalikes and inanimates (vs. animates) emerged in a cluster of central electrodes at 411 ms, compatible with the latency of ERP responses to contextual novelty at the level of categories (Campanella et al., 2000; 2002; Halgren & Marinkovic, 1995). Finally, to relate the present work to prior work (Bracci et al., 2019), we explored the ERP effects in a posterior cluster of electrodes, more closely capturing occipitotemporal activity. Consistent with the effect in the central cluster, here we found that ERPs differed between animates and lookalikes presented in the same context (inanimate-standards), but not between inanimates and lookalikes presented in the same context (animate-standards). This pattern emerged earlier in the posterior cluster than in the central cluster (from 227 ms vs. 411 ms post-stimulus onset), within the temporal window typically associated with visual object categorization (≈150–300 ms; Carlson et al., 2013; Cichy et al., 2014; Contini et al., 2017). This posterior-to-central temporal progression is consistent with a hierarchical organization of visual processing, in which categorical information is first computed in occipitotemporal cortex and subsequently propagates to more anterior regions (Cichy et al., 2014; Kietzmann et al., 2019; Bankson et al., 2018). Finally, greater representational similarity between lookalikes and inanimates was also found in VGG-19, a computational model of occipitotemporal cortex (Khaligh-Razavi & Kriegeskorte, 2014; Cichy et al., 2016).

Altogether, converging measures of biological and artificial object perception show that attribution of animacy overrides superficial animal appearance in visual object categorization, yielding accurate animate-inanimate distinction also for objects that carry features proper of animals. The alignment between the DNN analysis and human neural data underscores the potential of DNNs as computational models of human perception.

The first result of our study is that all three categories––animate, inanimate and lookalike––were accurately discriminated and categorized: the EEG response at the oddball frequency was strongly significant in all four conditions with a widespread effect peaking in posterior (visual) areas (Figure 2A). This is consistent with previous fMRI results by Bracci et al. (2019), who showed category discriminability using neural response patterns extracted from the visual ventral occipitotemporal cortex (VTC).

Different from our results, however, Bracci et al. (2019) found that, particularly in the anterior VTC, object representation was primarily driven by animal-like appearance: representation of lookalikes was more similar to representation of animate, than inanimate objects. The apparent discrepancy may stem from differences in task design, stimuli and/or methodology. In our study, participants passively looked at the stimuli. Instead, in Bracci et al., they were instructed to perform two tasks during stimulus presentation: an animacy task (“Does this image depict a living animal?”) and an animal-appearance task (“Does this image look like an animal?”). Both tasks might have specifically emphasized the relationship––the visual similarity––between lookalikes and animals. And, in effect, neural data showed no differences across task sessions, in any of the performed analyses. Moreover, Bracci et al.’s study involved nine lookalike objects, whereas our study used a set of 72 exemplars that naturally spanned a broad range of animal resemblance (from 23.50 to 83.50 on a 1-100 Likert-scale). We found that the degree of animal resemblance systematically modulated the neural (ERP) response to oddball-lookalikes. However, we also found that high animal appearance did not change the category boundaries: lookalikes with the highest animal appearance were still represented as more similar to inanimate objects than to animate ones. In this light, even if Bracci et al.’s lookalikes had the highest degree of animal resemblance, we would still expect to observe greater representational similarity between these lookalikes and animate objects.

A more plausible explanation for the apparent discrepancy between our results and those of Bracci et al. (2019) could lie in the methodology, and the measures of categorization used in the two studies. On the one hand, fMRI may be more sensitive to localized effects, such as the representation of animal features in specific VTC areas, which would capture the visual similarity between animals and lookalikes. On the other hand, the categorization response measured with ftEEG may be better suited to capture explicit categorization, predicting how an individual consciously represents an object (Retter et al., 2020; Spriet et al., 2025). In our data, a trace of the proposed “animal bias” in visual cortex is reflected in the posterior occipitotemporal correlations with animal-like appearance (Figure 4B). These correlations showed that, among inanimate objects, lookalikes with high animal-like appearance *looked odder* inducing greater oddball ERP amplitudes; whereas, among animate objects, lookalikes with lower animal-like appearance *looked odder* inducing lower oddball ERP amplitudes.

However, our results show that, if an animal bias exists in VTC, it is readily overcome, possibly through the integration of other ‘inanimate’ features, yielding truthful classification of lookalikes as inanimate objects. In this perspective, the animal bias reported in VTC would reflect great visual sensitivity to animal-like features rather than category misclassification in visual perception. Supporting this interpretation, Bracci et al. showed that participants were not misled by animal-like appearance and consistently judged lookalikes as inanimate objects.

Returning to our initial question: what is animacy in visual perception, and what does it *look like*? A set of studies have shown that animacy perception correlates with a range of visual features from spatial frequency spectrum to color, texture and shape (e.g., Jozwik, Kriegeskorte, Mur, 2016; Long et al., 2018; Nasr et al., 2014; Proklova, Kaiser, Peelen, 2016; Spriet et al., 2025) as well as with higher-level properties such as, for instance, agency and predictability (Gobbini et al., 2011; Jozwik et al., 2022). It also correlates with the presence of animal-like features, including faces, eyes and limbs (Bracci et al., 2019; Chen et al., 2023; Ritchie et al., 2021; but see Jozwik et al., 2023). However, our results suggests that animacy cannot be reduced to the property of having eyes, face, body or body parts; and perhaps it cannot be reduced to any single feature at all. In line with this, a recent study (Leys et al., 2025) using multivariate EEG analyses found distinct temporal windows for face/body perception (with peak around 150 ms) and for object discrimination by animacy (300-500 ms).

Considering the broad literature in the field, animacy perception may be conceptualized as a case of *lack-of-invariance problem* in vision. The lack-of-invariance problem refers to the phenomenon in speech perception whereby the categorization of a sound (e.g., /t/) is not determined by any single acoustic feature but instead depends on the surrounding sounds in which the sound is produced (Liberman et al., 1967; Hochmann & Papeo, 2014). By analogy, visual representation of animacy may result from the combination of a range of visual and non-visual features, at different level of complexity. This framework would explain why, given certain motion signals, individuals can perceive abstract shapes (squares and circles) as animate agents (Varrier, Su, et al., 2025; Wheatley et al., 2007), and how, as shown here, we are not confused when we see an object looking like a pig, on a shelf.

An interesting open question concerns the developmental trajectory through which human adults become accurate and efficient at discriminating between animate and inanimate entities, even in case of ambiguity. It has not been systematically investigated whether infants initially show a bias to identify ambiguous objects as animals or animates, before learning corrects this bias. However, there is evidence that infants as young as 4 months can categorize unfamiliar objects by animacy, suggesting an early capacity to exploit visual information in the input for object recognition (Spriet et al., 2022; Spriet et al., 2025; Peykarjou, Hoehl, Pauen, 2024; Xie et al., 2022). Another interesting avenue for future research is the study of other behaviorally relevant cases of visual ambiguities, such as robotic agents, food that resembles non-edible material or non-edible material that looks like food. Such studies could help probe the limits of human visual perception in other object domains (Dehn et al., 2025). Finally, while the present DNN analysis demonstrated convergence between artificial and biological vision, the specific computational mechanisms underlying this convergence remain to be understood (Duyck et al., 2024).

In conclusion, the investigation of lookalike object perception addressed the fundamental question of whether human visual perception is characterized by an animal bias, such that *anything* with eyes, face and/or body parts is *seen as* an animate, or it readily reconciles animal appearance with other, more veridical categorical information. Our findings support the latter interpretation: although animal-like appearance modulates the neural response to ambiguous objects, it does not override the categorical boundary between what is *alive* and what it is not. Animal-like features of objects are represented in visual cortex but their presence or absence does not constitute a critical dimension for visual object categorization by animacy.

## Data availability

Stimuli, EEG data and code for the main analyses will be deposited in the Open Science Framework repository created for this project.

## Authors’ contributions – CrediT

*Farzad Rostami*: Methodology, Formal Analysis, Investigation, Writing – Original Draft, Writing – Review & Editing, Visualization; *Céline Spriet*: Conceptualization, Formal Analysis, Investigation, Data Curation, Writing – Review & Editing, Visualization; *Hans Op de Beeck*: Conceptualization, Methodology, Writing – Review & Editing. *Jean-Rémy Hochmann*: Conceptualization, Methodology, Writing – Review & Editing, Supervision; *Liuba Papeo*: Conceptualization, Methodology, Writing – Original Draft, Writing – Review & Editing, Supervision, Funding Acquisition.

## Acknowledgments

We thank Karina Cazali for helping with data collection. L. P. was supported by an ANR-DFG Franco-German Grant (FRAL_RELATIONS_268980). F.R. was supported by a fellowship of the LabEx CORTEX of the University of Lyon (ANR-11-LABX-0042). C.S. was supported by a fellowship of the Fondation pour la Recherche Médicale (FDT202304016547).

